# Progress Towards a Public Chemogenomic Set for Protein Kinases and a Call for Contributions

**DOI:** 10.1101/104711

**Authors:** David H. Drewry, Carrow I. Wells, David M. Andrews, Richard Angell, Hassan Al-Ali, Alison D. Axtman, Stephen J. Capuzzi, Jonathan M. Elkins, Peter Ettmayer, Mathias Frederiksen, Opher Gileadi, Nathanael Gray, Alice Hooper, Stefan Knapp, Stefan Laufer, Ulrich Luecking, Susanne Müller, Eugene Muratov, R. Aldrin Denny, Kumar S. Saikatendu, Daniel K. Treiber, William J. Zuercher, Timothy M. Willson

**Affiliations:** Structural Genomics Consortium, UNC Eshelman School of Pharmacy, University of North Carolina at Chapel Hill, NC 27599, USA; AstraZeneca, Darwin Building, Cambridge Science Park, Cambridge, CB4 0FZ, UK; Drug Discovery Group, Translational Research Office, University College London School of Pharmacy, 29-39 Brunswick Square, London, WC1N 1AX; Miami Project to Cure Paralysis, University of Miami Miller School of Medicine, Miami, FL-33136, USA; Peggy and Harold Katz Family Drug Discovery Center, University of Miami Miller School of Medicine, Miami, FL, 33136, USA; Laboratory for Molecular Modeling, Division of Chemical Biology and Medicinal Chemistry, UNC Eshelman School of Pharmacy, University of North Carolina at Chapel Hill, NC, 27599, USA; Structural Genomics Consortium, Universidade Estadual de Campinas - UNICAMP, Campinas, SP, Brazil; Boehringer Ingelheim RCV GmbH & Co KG, Vienna 1121, Austria; Novartis Institutes for BioMedical Research, Novartis Campus, Basel, CH-4056, Switzerland; Structural Genomics Consortium and Target Discovery Institute, Nuffield Department of Clinical Medicine, University of Oxford, Old Road Campus Research Building, Oxford, OX3 7DQ, UK; Harvard Department of Biological Chemistry and Molecular Pharmacology, Harvard Medical School, Boston, Massachusetts 02115, United States; Department of Cancer Biology, Dana–Farber Cancer Institute, Boston, Massachusetts 02215, United States; Structural Genomics Consortium, Buchmann Institute for Molecular Life Sciences, and Institute of Pharmaceutical Chemistry, Goethe University Frankfurt, Max-von-Laue-Straβe 15, D-60438 Frankfurt am Main, DE; Department of Pharmaceutical Chemistry, Institute of Pharmaceutical Sciences, Eberhard Karls Universität Tübingen, Auf der Morgenstelle 8, 72076 Tübingen, Germany; Bayer Pharma AG, Drug Discovery, Müllerstrasse 178, 13353 Berlin, Germany; Worldwide Medicinal Chemistry, Pfizer Inc., Cambridge, MA 02139, USA; Global Research Externalization, Takeda California, Inc., 10410 Science Center Drive, San Diego, CA 92121, USA; DiscoverX Corporation, 42501 Albrae Street, Fremont, CA 94538

## Abstract

Protein kinases are highly tractable targets for drug discovery. However, the biological function and therapeutic potential of the majority of the 500+ human protein kinases remains unknown. We have developed physical and virtual collections of small molecule inhibitors, which we call chemogenomic sets, that are designed to inhibit the catalytic function of almost half the human protein kinases. In this manuscript we share our progress towards generation of a comprehensive kinase chemogenomic set (KCGS), release kinome profiling data of a large inhibitor set (Published Kinase Inhibitor Set 2 (PKIS2)), and outline a process through which the community can openly collaborate to create a KCGS that probes the full complement of human protein kinases.

## Kinases: Important targets, untapped opportunities

Protein kinases are a large family of enzymes that catalyze the transfer of phosphate from ATP to serine, threonine, and tyrosine residues of their substrate proteins. Protein kinases are found in all eukaryotes from yeast to mammals. Humans express 518 protein kinase catalytic domains using the Manning classification.^1^ The precise number, however, is a matter of debate, since a number of the proteins are pseudokinases that lack catalytic activity, some have been re-characterized as members of other protein families (e.g. bromodomains) and some are enzymes for non-protein substrates.

Over the course of 20 years of experimentation medicinal chemists have become adept at synthesizing cell-active kinase inhibitors that target the ATP binding site in the catalytic domain of these enzymes.^2–4^ Coupled with the discovery that protein kinases are involved in almost every aspect of cell signaling and are often dysregulated in human diseases, these enzymes have become popular targets for development of drugs. Academic, biotech, and large pharmaceutical company labs alike have pursued kinase inhibition creatively and diligently. These efforts have borne fruit, and the FDA has approved over 35 small molecule kinase inhibitor medicines since the turn of century. Accordingly, protein kinases have proven to be among the most productive of human gene families for development of targeted therapeutics.^5–6^

Despite the success of protein kinase drug discovery within the pharmaceutical industry, much of their therapeutic potential remains untapped. An analysis of peer-reviewed publications and published patent applications in 2010 revealed that 80% of the protein kinases remained poorly studied and their roles in human biology were largely undefined.^7^ Moreover, the vast majority of kinase medicinal chemistry and experimental data remains firewalled within company databases or held as proprietary know-how, which means the broader scientific community has limited access to this body of knowledge. We anticipate that a wide range of human diseases will be amenable to treatment with inhibitors of the currently untargeted protein kinases and that if a high quality set of tool molecules for all protein kinases were freely available, it would enable their role in cell signaling to be better understood.^8^ Building this detailed understanding of kinase roles on a genomic scale together as a community will contribute to the validation of useful kinase targets across the kinome. These identified targets can then be considered for the aggressive pursuit, commitment, and action required to bring new medicines to patients.

One way to understand the role of a particular kinase is to utilize a chemical probe for the kinase of interest. Small molecule chemical probes that meet stringent criteria for potency and selectivity are powerful tools to study the biology of their target proteins in cells.^9^ Several investigators have successfully identified chemical probes for historically understudied protein kinases. These probes proved useful to understand the role of the specifically targeted kinase in disease biology. For example, synthesis of a chemical probe for BRAF facilitated the study of its role in tumorigenesis and eventually led to the development of several anti-cancer drugs.^10^ Likewise, chemical probes for the poorly studied kinases such as ZAK^11^, STK16^12^, PLK4^13^, and STK33^14^ have advanced research in oncology and other diseases. However, although these probes proved extremely useful, each report also highlights the challenges in identification of highly selective ATP-competitive inhibitors, which can be resource intensive given the large number of kinases in the protein family.

Building a set of over 500 highly selective kinase chemical probes will take many, many years of concerted effort. In addition, the community does not know how to prioritize the development of individual chemical probes. In what order should we address the remaining kinome so that we find the best targets first? Is there another, more efficient way to identify kinase targets that are worth pursuing in more depth?

## Kinase Chemogenomics

To expedite kinome-wide target discovery, we have begun construction of a comprehensive kinase chemogenomic set (KCGS). This practical solution takes advantage of the chemical connectivity of kinases (cross reactivity of inhibitors), the large numbers of kinase inhibitors already made by labs around the world (and thus volume of data available), and the ability to screen practically kinome wide. We call the methodology “kinase chemogenomics” since it seeks to use a set of small molecule kinase inhibitors to interrogate the biology of all kinase gene products in cells. Bunnage and Jones recently reviewed the concept and utility of chemogenomic libraries.^15^ They describe a chemogenomic set as a collection of small molecules with defined (annotated) and narrow activity. A hit from the testing of such a library in a phenotypic screen implies that the annotated targets of that hit are involved in modulation of the measured phenotype. Our KCGS is being constructed with this exact goal in mind: hits from the set will point to potential targets driving phenotypic responses in disease relevant phenotypic assays. These targets then are candidates for follow up experiments such as target knockdown with CRISPR-Cas9 or RNAi, or chemical probe development (**Figure 1**). These more pointed biological studies can add further support to the utility of the identified targets. The successful interplay between annotated compound sets, phenotypic screening, genetic methods, and bioinformatics is well documented, and integration leads to new insights on targets and pathways of interest.^16–23^ Herein we describe our progress towards the generation of a publicly available KCGS, and outline a collaborative plan to complete the set. This collaborative project between industrial and academic scientists will build a comprehensive KCGS composed only of potent, narrow spectrum inhibitors that collectively demonstrate full coverage of all human protein kinases for which there are assays available at one of the contract research organizations that offers broad kinome profiling (“the screenable kinome”). A publicly available comprehensive KCGS will accelerate basic research into the biological function and therapeutic potential of hundreds of understudied protein kinases.

**Figure 1.**
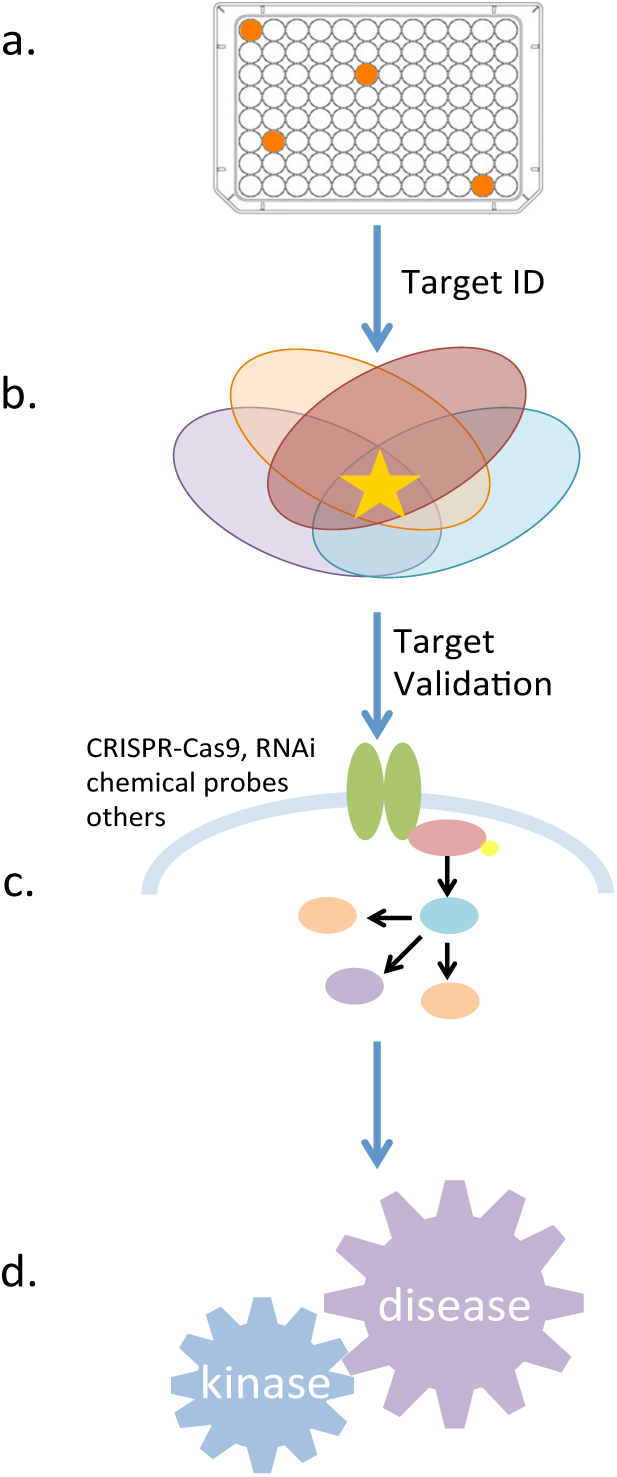
Schematic: Utilization of the KCGS. (**a**) Disease relevant phenotypic screen highlights active compounds (**b**) Because all the targets are annotated, active molecules points to potential targets of interest. (**c**) Follow up experiments can be used to help confirm importance of these highlighted targets (**d**) Body of evidence implicates kinases that can be inhibited to impact disease.

## The Published Kinase Inhibitor Set (PKIS): KCGS proof of concept experiments

We released our first kinase chemogenomic set, the Published Kinase Inhibitor Set (PKIS^24^), to catalyze new academic research on understudied kinases by provision of physical samples of a large set of kinase inhibitors, along with the annotation detailing kinase inhibitory profiles. PKIS is a collection of 367 ATP-competitive kinase inhibitors representing 31 diverse chemotypes.^25^ Design of the set capitalized on the fact that ATP-competitive inhibitors often demonstrate activity across multiple kinases. The inhibitors were chosen from molecules that had been previously published by medicinal chemists at GlaxoSmithKline, but had only been screened against a limited number of kinases. Screening of PKIS across 232 protein kinases identified potent inhibitors of kinase targets across the kinome, and showed that this set is made up of compounds with a range of selectivity profiles. The chemical structures of the inhibitors and their associated kinase screening data were placed in the public domain (https://www.ebi.ac.uk/chembldb/extra/PKIS/) and the set was made widely available to the research community. Using a web-based request mechanism (https://pharmacy.unc.edu/research/sgc-unc/request-pkis/) a physical copy of PKIS was distributed to over 200 academic investigators to support their research.

Sharing the PKIS compounds openly allowed investigators across the scientific community to query the impact of kinase inhibition in diverse biological contexts. For example, PKIS was used to identify potential medicinal chemistry starting points and generate ligand bound crystal structures for the understudied GRK family kinases.^26^ The GRKs have been implicated in heart failure^27^, Parkinson’s disease^28^, and multiple myeloma^29^. A PKIS compound was identified as a potent ligand for the pseudokinase MLKL, and the authors demonstrated that binding to this pseudokinase inhibited necroptosis.^30^ Fifty-three PKIS compounds were identified with sub-micromolar potency against *T. brucei*, the protozoan pathogen that causes human African trypanosomiasis, a devastating disease in the developing world.^31^ PKIS was used in a high content screen investigating axon growth in primary CNS neurons.^32^ The team was able to use the broad inhibitor annotation and phenotypic readouts to develop a semi-empirical, machine learning algorithm that allowed for generation of testable hypotheses as to patterns of kinase inhibition that led to neurite outgrowth. Screening of PKIS in a number of chordoma cell lines identified several different EGFR inhibitors with activity.^33^ Chordoma is a rare bone cancer, and no targeted therapies have been approved for use in chordoma patients. Identification of a well-studied kinase such as EGFR with drugs on the market and a number of compounds advancing in the clinic for other cancer indications, allows for potential repurposing, a faster way to approval. These and other examples^34–44^ of PKIS applications demonstrate the utility of sharing an annotated kinase inhibitor set and convinced us to proceed with the construction of a comprehensive KCGS – one that provides coverage of the currently screenable kinome.

## The Screenable Human Kinome

As an important step in building a comprehensive KCGS we sought to define the readily screenable human kinome—those protein kinases for which a robust assay is available through a commercial vendor. We only counted assays for wild type human protein kinases and excluded assays for mutant protein kinases, lipid kinases and sugar kinases (Supplemental Table 1). The ten commercial vendors surveyed were Carna Biosciences, DiscoverX, Eurofins, Luceome, MRC PPU, Nanosyn, ProQinase, Reaction Biology Corp., SignalChem and ThermoFisher. The commercial protein kinase assays are run in multiple formats, but all have been widely used by the kinase research community to measure kinase inhibitor activity and selectivity across a wide range of human kinases.^45^ Of the total kinome, we identified 436 kinases for which there were commercially available assays. Many of the kinases for which no assay was available have been classified as pseudokinases^46^ (**Figure 2a**). Of course, the absence of a contract research organization assay for a particular kinase does not mean that the kinase in question is not assayable, and in fact there are assays for some pseudokinases. Non-overlapping assays from only two vendors (**Figure 2b**) provides efficient profiling of the candidate inhibitors against 97% of the screenable human kinome. Importantly, for more than half of the screenable kinome (260 kinases), one can choose from up to seven different vendors, and this offers flexibility in terms of assay format, custom panel options, data visualization tools, and practical considerations such as turnaround time and pricing structure.

**Figure 2.**
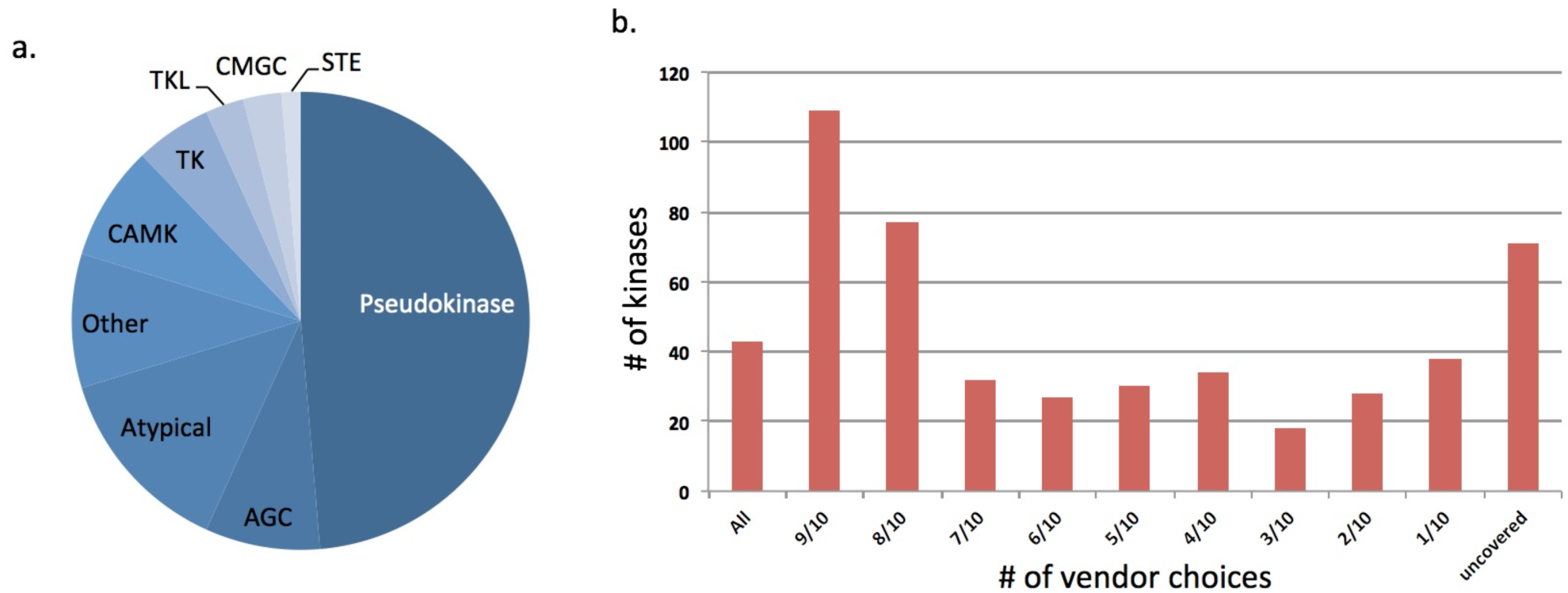
The screenable protein kinome. Using assays from 10 vendors a total of 436 unique non-mutant human protein kinases can be readily screened. (**a**) Subfamily representation of the protein kinases that were not available through the 10 vendors. Nearly half of the unscreenable human kinome is composed of pseudokinases (**b**) Histogram illustrating how many vendors can be used to screen across the kinome. For example, there are 43 kinases that all 10 vendors have screens for (“all” bar). There are another 109 kinases screened by 9 out of 10 vendors (“9/10” bar). There are 38 kinases that only 1 out of the 10 vendors has assays for (“1/10” bar). *Note: Our definition of the “screenable kinome” is those kinases for which there is a commercial assay that can be accessed. It is likely that assays could indeed be configured for many of the kinases not currently on this list.*

## Progress towards a comprehensive KCGS

Our ideal KCGS will contain 1000–1500 compounds. This number of compounds is on the order of what many disease relevant phenotypic screens can handle efficiently. In addition, we conducted a simulation (see methods section for further detail) to estimate the theoretical number of compounds required to achieve kinome coverage under the selectivity criteria set forth herein. The simulation suggested that on the order of 570 compounds was sufficient to achieve a set with all the desired attributes. Thus 1000–1500 compounds well-selected compounds can meet our goal of full kinome coverage and will be able to provide at least two to four distinct chemotypes (structural classes) inhibiting each kinase.

In kinase medicinal chemistry, different chemotypes that target the same kinase often have different off-target kinase profiles. When two or more chemotypes that inhibit a given kinase are active in a phenotypic screen, one gains confidence that the phenotypic activity is linked to that target kinase. Ideally, the compounds included in the KCGS will potently inhibit at least one kinase with an IC_50_ value less than 100 nM, and each compound in the set will have potent activity on less than 4% of the kinome. We will avoid inclusion of even moderately promiscuous compounds (inhibiting >10% of the screenable kinome). Promiscuous compounds do not aid in the deconvolution (target identification) of phenotypic screening results. Experimental results from a set of narrow spectrum kinase inhibitors, however, can help scientists focus on the kinase or sets of kinases that are driving the phenotypic response of interest. The comprehensive KCGS will be built from PKIS compounds, PKIS2 compounds (described below), appropriate literature compounds, and donations from consortium members. In the following section we describe each of these pieces in turn.

### PKIS compounds suitable for inclusion

As mentioned above, PKIS was developed as a first generation, proof-of-concept KCGS. PKIS compounds have a range of selectivity profiles, with some members of the set being broad spectrum, thus too promiscuous for inclusion in our KCGS. Because of this, we have defined criteria for a narrow spectrum inhibitor to be suitable as a member of the comprehensive KCGS (Box 1, and more detail in the methods section). Based on current data, 131 compounds from PKIS have useful potency and narrow spectrum inhibition profiles, and these compounds represent 22 different chemotypes and cover a total of 58 kinases (Supplementary Table 2).

##### Box 1. Criteria for kinase chemogenomic set inclusion

**PKIS:**

**Potency:** At least 1 kinase inhibited >90% Inhibi? on *and* one of these selectivity criteria must also be met:

> **GINI**: > 0.75
>
> **Entropy**: < 1.65
>
> **Selectivity index:** SI(90) < 0.02
>
> *\*If only GINI is met then the inhibitor must not inhibit more than 7 kinases above 90%I for KCGS set inclusion*

**PKIS2:**

**Potency** – At least 1 kinase with K_d_ < 100nM **Selec3vity index** – SI(65) < 0.04

### PKIS2 compounds suitable for inclusion

Building on the success of PKIS, we assembled a second kinase chemogenomic set (PKIS2), composed of 645 small molecule inhibitors representing 86 diverse chemotypes that were published by medicinal chemists at GlaxoSmithKline, Pfizer, and Takeda. PKIS and PKIS2 have only nine chemotypes in common. The PKIS2 compounds that are members of these nine chemotypes are not duplicates of PKIS compounds, but rather expand on the structural variety within the chemotype. Herein we report the composition of PKIS2 and broad kinome screening data for these compounds. We profiled PKIS2 in singlicate at a concentration of 1 µM against a broad panel of 392 wild-type human kinases using competitive displacement of diverse immobilized inhibitors as a measure of the activity of a compound on each specific kinase.^47^ The resulting kinome-wide profiling data showed tight distribution of highly selective compounds (**Figure 3**). 357 compounds that demonstrated significant activity on <4% of human wild type kinases were selected for confirmatory *K*_d_ determinations on 339 kinases. Specifically, we generated K_d_ values for all compounds with SI(65) < 0.04 against those kinases that showed >80% I. The K_d_ values can be found in Supplementary Table 3. The high confirmation rate of the *K*_d_ determinations demonstrated that the initial broad kinase screen had <0.5% false positive rate despite being run in singlicate.

**Figure 3.**
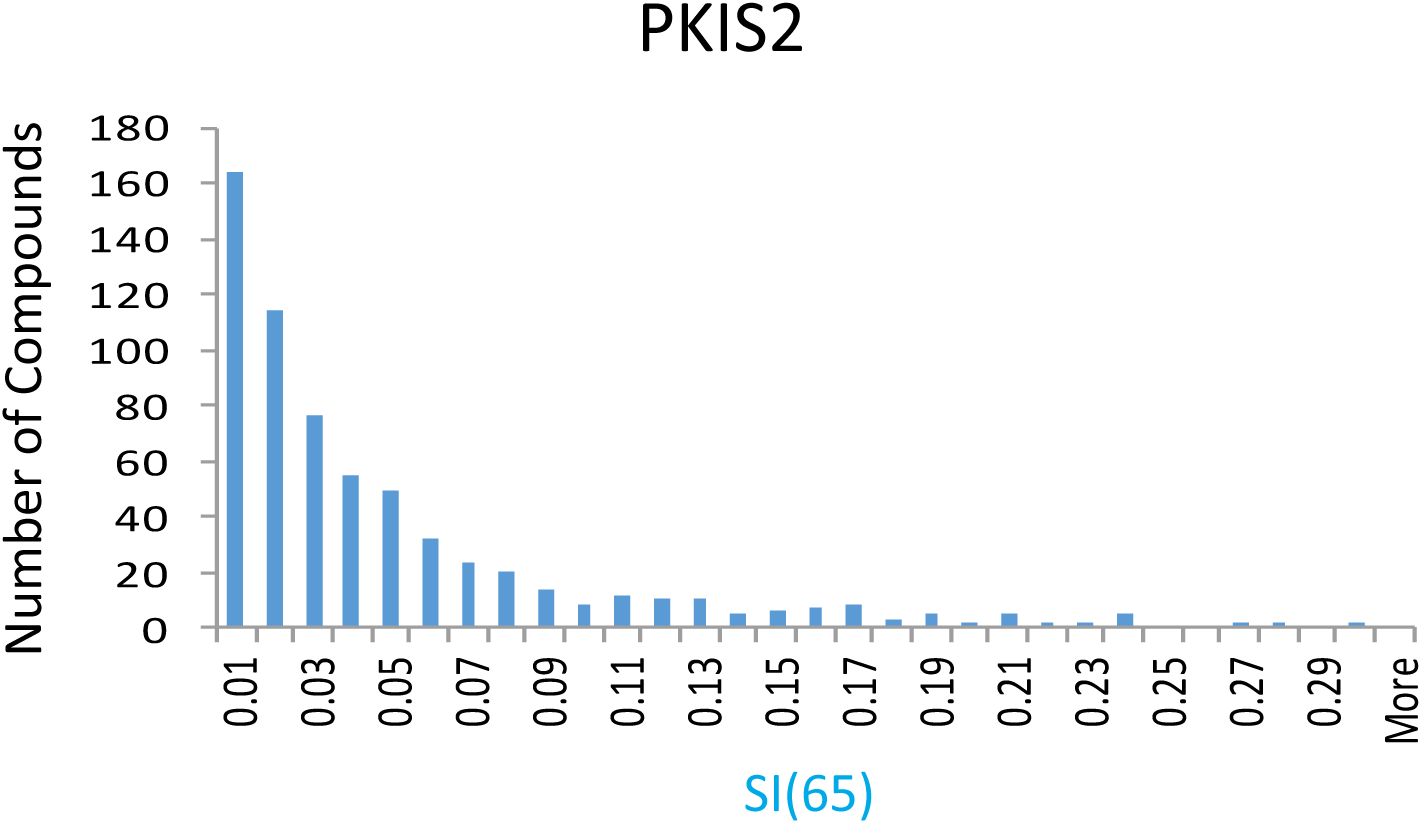
Selectivity profile of PKIS2 compounds using DiscoverX KINOMEscan. Selectivity Index analysis of PKIS2. 357 compounds demonstrate an SI(65) of <0.04 at 1 µM, and thus inhibit less than 4% of the kinases in this screening panel with more than 65% inhibition at the 1 μM screening concentration.

In addition, 90% of the confirmed actives demonstrated a *K*_d_ <1 µM at one or more kinases. All chemical structures and wild type kinase profiling data are available in Supplementary Table 4. Applying our potency and selectivity criteria for inclusion in the comprehensive KCGS, 174 PKIS2 compounds (Supplementary Table 5) from 23 chemotypes have narrow spectrum activity on 81 kinases.

Based on this analysis of the screening data we have for PKIS and PKIS2 compounds, these two sets contain potent and narrow spectrum compounds for 122 non-overlapping kinases. This represents coverage of 28% (122/436) of the screenable kinome.

### Literature compounds suitable for inclusion

With a goal of complementing PKIS/PKIS2, we surveyed the peer-reviewed literature for kinase inhibitors with activity on the existing gap kinases. This search was done in a non-automated manner, and thus it is likely there are high quality compounds we have not identified yet. Inhibitors were selected if the literature reported potency <100 nM activity on a gap kinase and if the compound appeared to display a suitable narrow spectrum profile through screening against at least 20 (ideally many more) other protein kinases. The selectivity element of this literature analysis is complicated by the wide variety of screening strategies utilized, with variation in compound screening concentration, composition of assay panel, assay format, and amount and detail of profiling data actually shared in each publication. Our analysis to date of publications originating from academia and industry has identified 152 potent and selective kinase inhibitors (Supplementary Table 6) that have activity on an additional 132 protein kinases (Supplementary Table 7) not covered by inhibitors selected from PKIS/PKIS2. These kinase inhibitors are candidates for inclusion into the KCGS pending permission to incorporate them, broader screening to standardize selectivity data, and confirmation of a narrow spectrum selectivity profile when screened across the kinome.

## A Virtual Kinase Chemogenomic Set

**Figure 4** provides examples of compounds from PKIS, PKIS2, and the literature that meet criteria for inclusion in the KCGS, and examples of compounds that do not warrant inclusion because of insufficient selectivity.

**Figure 4.**
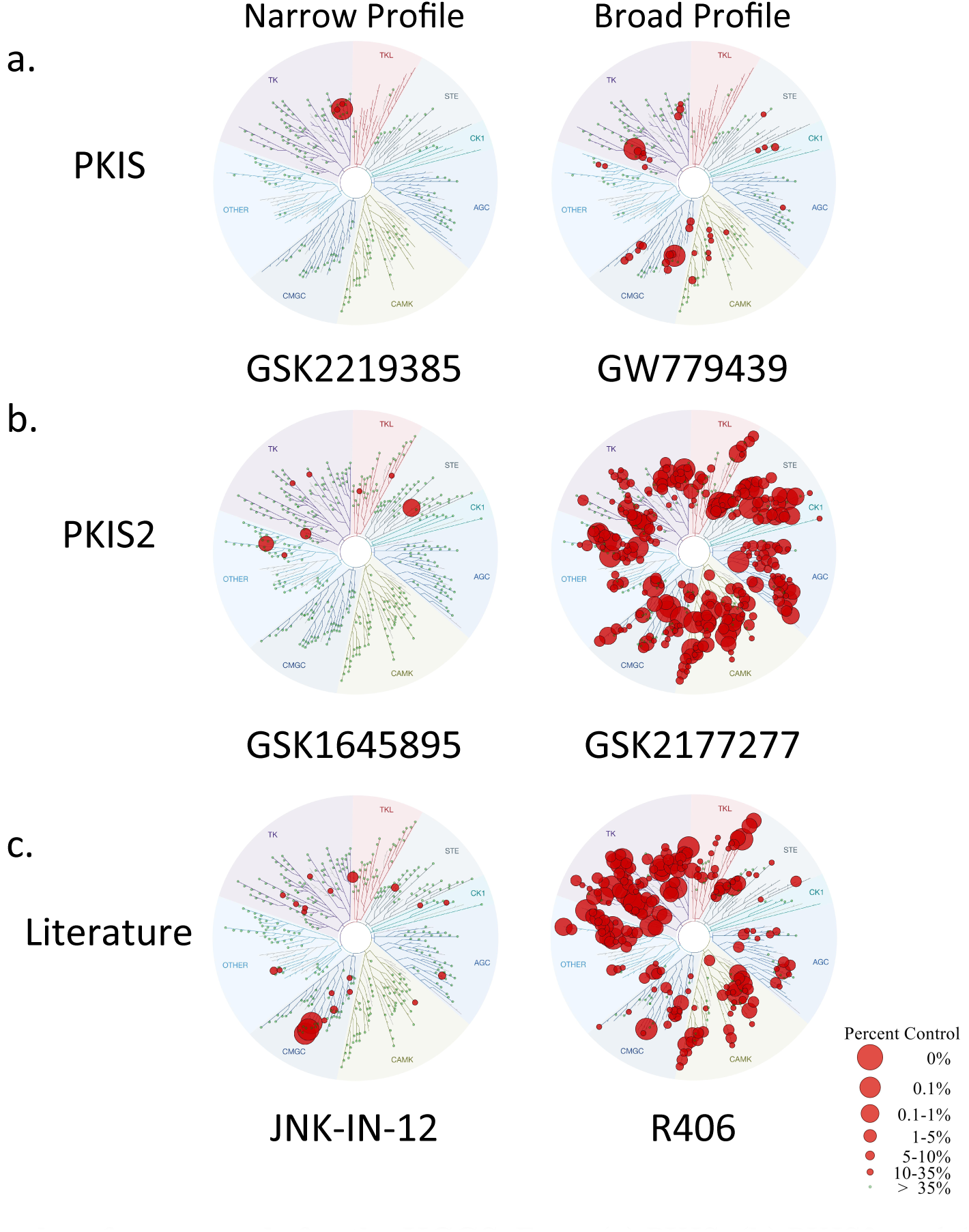
Examples of compounds for the KCGS. From (**a**) PKIS, (**b**) PKIS2, and the (**c**) literature that are suitable (narrow) or not suitable (broad) for inclusion in the KCGS.

In total we have identified 390 potential narrow spectrum inhibitors from PKIS, PKIS2, and the peer-reviewed literature that demonstrate activity on 254/436 of the screenable human protein kinases (**Figure 5**). Assembly of these compounds into a single chemogenomic set would yield a collection of inhibitors that have useful activity on 58% of the currently screenable protein kinome.

**Figure 5.**
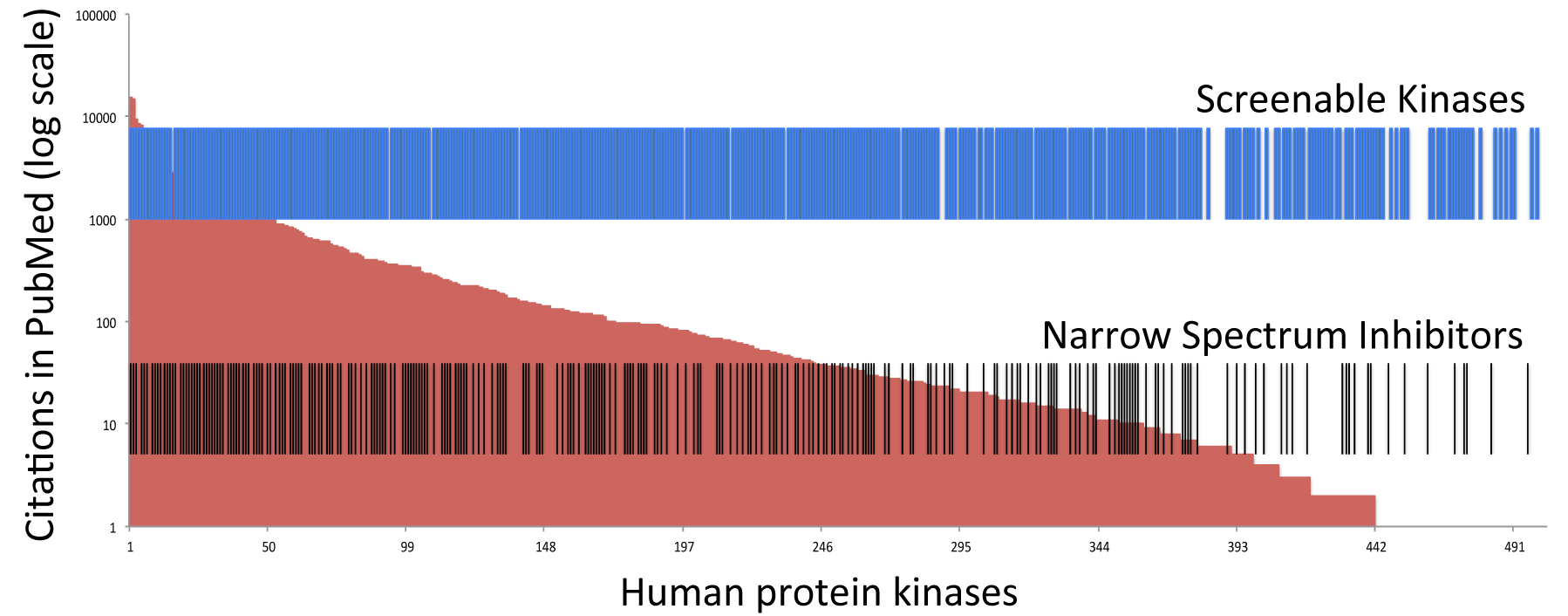
Representation of the kinome coverage of the virtual set of 457 narrow spectrum inhibitors. Red background shows the human protein kinases ranked by number of citations. Blue bars show the protein kinases for which an assay is available at one of 10 commercial vendors. Black bars identify protein kinases that are covered by the virtual set of inhibitors (subset of PKIS, PKIS2, and literature) described in the text.

However, there are limitations to this ‘virtual set’ of candidate chemogenomic compounds. First, many of the compounds have been screened on less than half of the currently available assays and may not meet our criteria for inclusion in the final set upon broader kinase activity profiling. Second, some of the published inhibitors may no longer be available in sufficient quantities for distribution requiring additional chemistry to resynthesize the compounds. Third, the set still fails to address 42% of the screenable protein kinome. Thus, additional work is required to reach the goal of building a comprehensive KCGS.

## The Kinase Chemogenomic Consortium

To complete the design and construction of a KCGS with complete coverage of the screenable protein kinome, we have assembled a consortium of academic and industrial partners. The consortium has the following operating principles:

1. The comprehensive KCGS will be created as an openly available public resource to support basic research on kinases.
2. All compounds in the set will be narrow spectrum inhibitors meeting the potency and selectivity criteria defined above.
3. Kinase profiling will be performed using the currently accessible commercial assays, and the set will be profiled in new assay formats that offer additional information as these become available (for example, in cell target engagement).
4. All chemical structures and kinase activity data will be made publically available.
5. The comprehensive KCGS will be made available to all scientists that agree to be trustees of the set.^48^

The current PKIS/PKIS2/literature virtual set serves as the starting point for development of the comprehensive KCGS. We started this endeavor in 2016 and asked pharmaceutical company and academic partners for their ideas and input on set generation (**Figure 6**).

**Figure 6.**
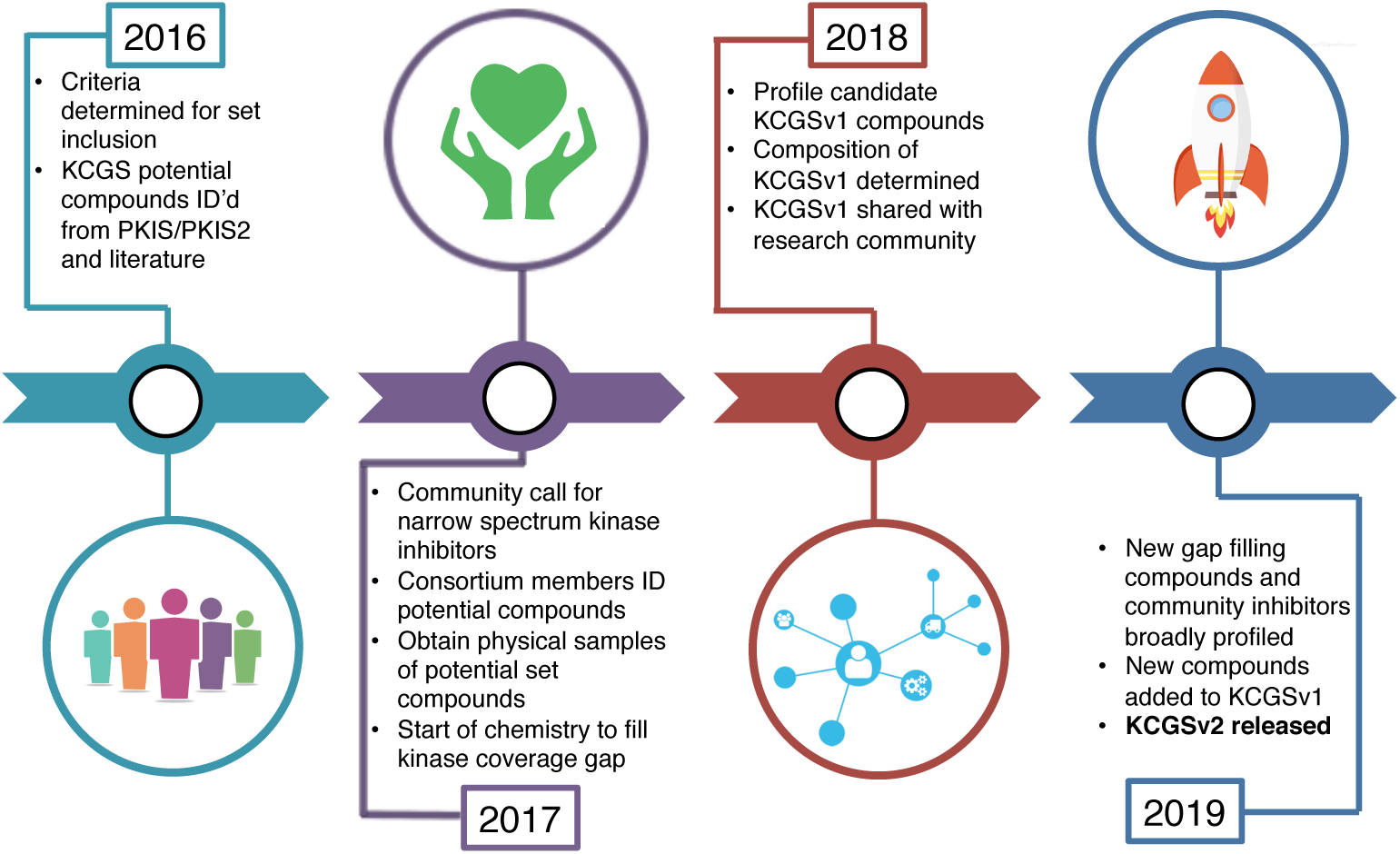
Process map for construction of a public comprehensive protein kinase chemogenomic set (KCGS).

GSK, Pfizer and Takeda were the first members of the consortium to physically contribute samples and we anticipate several other pharmaceutical companies and academic labs will contribute to the set in 2017. This year, to start filling the gaps (gap kinases listed in Supplementary Table 8), we have asked current consortium members to identify and provide physical samples of kinase inhibitors that will cover additional kinases and likely have the requisite potency and selectivity to be included in the set. Our intent is that this publication serves as a call to *all* contributors in the research community to suggest other possible candidates from the literature or their own research to supplement this work. To be considered for the set the compounds must meet minimum criteria:

- Potency – more potent than 100 nM on gap kinase of interest
- Selectivity – profiled against at least 100 kinases with no more than 5% of the screened kinases more potent than 100 nM or >90% inhibition at 1µM

The compounds that we receive physical samples of will be sent for profiling against the screenable kinome. This will enable direct comparison of compounds in the same set of assays and conditions to evaluate potency and selectivity, and provide the information needed to make decisions on what compounds should be included in the final physical set. This data will also be placed in the public domain.

We realize that inhibitors of suitable quality may not currently exist for some of the uncovered kinases. For example, inhibitors that do exist for these gap kinases may be too promiscuous or not potent enough. This paper is also a call for input from the kinase medicinal chemistry community on what molecules to synthesize for these uncovered kinases based on available data and insight. If you can recommend modest starting points for gap kinases (potency between 100-500 nM; active on 5-15% of the kinome; ideally a chemotype not represented in PKIS/PKIS2), we would welcome a collaborative effort in which we work together on synthesis, screening, and funding of the project.

Current members of the consortium include the academic and industrial laboratories represented as coauthors of this paper. More importantly, membership of the kinase chemogenomic consortium remains open to any scientist who wishes to contribute to the creation of the comprehensive KCGS as a public resource. Membership in the consortium allows for access to pre-publication kinase screening data, medicinal chemistry collaboration with our team, broad kinome screening as required to ensure compound suitability, and a copy of the complete set, once finalized and created, for screening in phenotypic screens. In summary, we welcome contributions of existing kinase inhibitors and submission of ideas for synthesis of new inhibitors that address any of the gap kinases. Scientists with interest in joining the consortium should contact David Drewry by email at.

## Conclusion

Creation of a comprehensive protein kinase chemogenomic set is a large undertaking that can best be accomplished as a collaborative effort. Importantly, notwithstanding the scale of the endeavor, it is a tractable problem. The combined effort of medicinal chemists over 20 years of kinase research has already laid the foundation to the extent that many of the compounds and much of the data already exist. While no one research group, university, or commercial organization has access to all of the best inhibitors, together as a community we can pool resources and knowledge to achieve the goal. Many of the compounds have already been synthesized, most of the screening assays are commercially available, and importantly there is strong scientific rationale to explore the full human kinome.

The comprehensive KCGS will have it greatest value only if it is freely available as a public resource. Use of the set in diverse disease relevant phenotypic screens and sharing of the resulting data in the public domain is the best mechanism to ensure that the therapeutic potential of as many protein kinases as possible will be uncovered. This effort will bring to light those kinases that deserve more detailed drug discovery efforts. Finally, additional incentive to build this set is provided by the important roles protein kinases have in non-mammalian species. Although the comprehensive protein kinase chemogenomic set will be designed to explore the human protein kinases, cross-species homology makes it likely that many of the inhibitors will have useful activity on parasite^49^ and plant kinases^50–51^. Thus, the work of the consortium will provide tools for other research sectors, and may lay the groundwork for development of comprehensive protein kinase chemogenomic sets for these other species.

Looking forward, the opportunity to use chemogenomic sets to explore the biology of other protein families beyond kinases cannot be overlooked. Analysis of many other druggable protein families reveals additional examples of the Harlow-Knapp effect: 90% of the research is focused on only 10% of the proteins^52^. For protein families where chemical connectivity between inhibitors can be demonstrated, such as phosphodiesterases (PDEs)^53^, histone deacetylases (HDACs)^54^, bromodomains^55^, and non-olfactory G-protein coupled receptors (GPCRs)^56^, we propose that development of publically available chemogenomic sets will be a profitable approach to expand our biological understanding of the human genome.

## Methods

### Simulations to estimate adequate set size for full kinome coverage

We conducted a simulation to estimate the theoretical number of compounds required to achieve kinome coverage under the selectivity criteria set forth herein. To accomplish this, we integrated over 1 million compound-kinase activity data points collected from the literature (examples^57–61^) and chemical repositories (PubChem, ChEMBL). We used these data to calculate probabilities of co-inhibition at ≥ 1 µM potency for pairs of kinases that were represented in the aggregate dataset. A total of 427 kinases had sufficient profiling data for inclusion in this exercise. The simulation of a compound’s activity profile began by assigning activity to a randomly selected kinase, following which the activity was “diffused” to a variable number of additional kinases. Activity diffusion was constrained by the calculated probabilities of co-inhibition in order to reflect the pharmacological linkages observed between kinases within the existing chemical space. Simulated inhibition profiles that targeted ≤ 15 kinases at the computed potency level were appended to the virtual set, provided they did not overlap with previously incorporated profiles by more than 3 targets, and they did not include targets that are already inhibited by 5% of the total set of simulated compound profiles. Repeating this sequence until every kinase was inhibited by at least 3 simulated compounds revealed that an average of about 571 compounds was sufficient to achieve a set with all the desired attributes. Increasing the number of kinases from 427 to about 500 will require increasing the number of compounds to maintain coverage criteria. However, our current results suggest that this requirement is likely to remain well within the 1000-1500 compound limit.

### PKIS2 Compound Preparation and Screening in KINOME*scan*

All PKIS2 compounds were dissolved and stored at -20°C as 10mM DMSO stocks. PKIS2 screening was performed using the commercially available KINOME*scan* assay panel as previously described.^47^ These assays are based upon a competition-binding assay that measures the ability of a small molecule to compete with an immobilized inhibitor for the active site. We screened the set at 1 μM in singlicate against 468 kinases (including mutants and non-human kinases). The results are provided as the percentage of the kinase that remains bound to the immobilized inhibitor relative to a DMSO control (% control). We then converted these numbers into percent inhibition (%Inh):

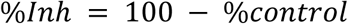

Since the assay was only run in singlicate it was important to follow up actives with K_d_ determinations. We determined which compounds would get K_d_ values determined based on the SI(65), which is the number of kinases bound <65% divided by the total number of assays. The compounds that had an SI(65) < 0.04 and at least 1 kinase inhibited above 90% were followed up by obtaining K_d_ values for any kinase that was inhibited >80%. Any kinase inhibitor that had an SI(65) less than 0.04, and was active ( < 100nM) on at least 1 kinase in these follow up K_d_ experiments, was marked as being a candidate for the KCGS.

### PKIS Compound Selection into the Virtual Kinase Chemogenomic Set

PKIS is a set of 367 kinase inhibitors originally profiled against 220 kinases at Nanosyn at 100 nM and 1000 nM.^25^ For evaluation of compounds to be included in the chemogenomic set activity data at 1000 nM was used.

Selectivity Index (SI): For each compound, the SI was assessed at a threshold of 90% kinase inhibition. The SI was then computed as:

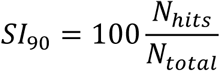

In this equation, SI_90_ is the selectivity index at 90% inhibition, N_hits_ is the number of kinases inhibited by the compound at ≥90%, and N_total_ is the total of kinases against which the compound was tested. It should be noted that the selection of the inhibition threshold is arbitrary and will influence the perceived selectivity of the compound. Moreover, kinase panels of different sizes should not be directly compared, as the SI is predicated upon N_total_.

Gini Coefficient: The previously described methodology for the use of the Gini coefficient to express kinase inhibitor selectivity was followed.^62^ In brief, this application of the Gini coefficient plots the cumulative fraction of total inhibition against the cumulative fraction of assayed kinases, arranged from least inhibited to most inhibited, and calculates the ratio of the area between the linear diagonal and cumulative inhibition curve to the area under the linear diagonal. A linear increase in cumulative inhibition thus results in a value closer to zero and indicates low selectivity. Compounds with high selectivity, on the other hand, show a slow rise in cumulative inhibition at the beginning, and a steep rise towards the end (Lorenz curve), thereby resulting in a Gini coefficient closer to 1. Values closer to 1 indicate “selective” compounds, while values approaching 0 indicate “promiscuous” compounds. It should be noted that Gini coefficients can be computed regardless of the number of kinases against which the compound was tested. Thus, the values for any compound can be directly compared.

Entropy Score: The entropy score, previously described by a team at the Netherlands Translational Research Center to identify selective kinase inhibitors quantifies the distribution of a compound’s inhibition activity over a panel of kinases, analogous to the distribution of a thermodynamic system over multiple energy states.^63^ Thus a small entropy score indicates high selectivity (narrow distribution), while a high entropy score indicates low selectivity (broad distribution). The score was originally developed for IC_50_ (or K_d_) measurements and was modified for single-point inhibition data. The following steps were implemented to calculate the entropy score for each compound within the panel of kinases against which it was tested:

1. Convert percent inhibition (%I) at 1000 nM (1 µM) to IC_50_ using the equation:

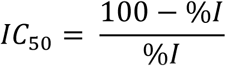

*note: if 100% inhibition was the reported result, 99.5%I was used in this calculation*
2. Compute K_*a*_ values for each kinase by 1**/**IC_50_
3. Sum all respective K_*a*_ values to generate ΣK_*a*_
4. For each individual K_*a*_ value, calculate K_*a*_**/** ΣK_*a*_
5. For each individual K_*a*_, then calculate (K_*a*_**/** ΣK_*a*_)*ln(K_*a*_**/** ΣK_*a*_)
6. Sum each associated value from (5) and multiple by -1 to obtain the resultant entropy score

Compounds from PKIS were selected as candidates for the KCGS if they met one of three criteria:

<u>Gini</u>: >0.75
<u>Entropy</u>: <1.65
<u>Selectivity index S(10) at 1 μM</u>: 0<x< 0.02
<u>Potency</u>: At least 1 kinase inhibited >90% Inhibition

If only the Gini measurement criterion was met then for that compound to be considered it had to inhibit no more than 7 kinases above 90% inhibition at the 1 μM screening concentration. For purposes of set inclusion only the Nanosyn data was utilized.

### Literature Compound Selection for the Kinase Chemogenomic Set

Due to the varying assay types and formats that were employed in the literature some basic rules were followed to identify compounds that have the potential to be included in the KCGS. For eventual inclusion, systematic broad screening will need to be completed.

Panel size – compound must have been screened against at least 20 other kinases to assess selectivity; if the panel is small the kinases represented should come from different parts of the kinome

Potency – must be at least 100 nM or 90% inhibition at 1 μM

Selectivity – must not inhibit more than 10 kinases above 90% inhibition at 1μM

## Acknowledgements

This work was funded by the UNC Eshelman Institute for Innovation.

The SGC is a registered charity (number 1097737) that receives funds from AbbVie, Bayer Pharma AG, Boehringer Ingelheim, Canada Foundation for Innovation, Eshelman Institute for Innovation, Genome Canada, Innovative Medicines Initiative (EU/EFPIA) [ULTRA-DD grant no. 115766], Janssen, Merck & Co., Novartis Pharma AG, Ontario Ministry of Economic Development and Innovation, Pfizer, São Paulo Research Foundation-FAPESP, Takeda, and Wellcome Trust [092809/Z/10/Z].

The authors thank Aled Edwards (SGC), Christian Fischer (Merck & Co.), Amy Donner (SGC and Chemical Probes Portal), Heather King (Chemical Probes Portal), Dave Morris (SGC and UNC Catalyst for Rare Diseases), and Matthew Robers (Promega Corp.) for helpful discussions and feedback.

GlaxoSmithKline, Pfizer, and Takeda are gratefully acknowledged for donation of PKIS2 compounds.

## Supplementary Information Available

Table 1 – Kinase assays by vendor

Table 2 – List of PKIS1 compounds that meet criteria for KCGS inclusion

Table 3 – PKIS2 K_d_ determinations

Table 4 – Full PKIS2 dataset from KINOME*scan* panel

Table 5 – List of PKIS2 compounds that meet criteria for KCGS inclusion

Table 6 – List of literature compounds that meet kinase potency and selectivity criteria but will require additional screening to ensure suitability for KCGS

Table 7 – List of additional kinases (beyond those covered by PKIS and PKIS2 compounds) that can be covered by literature compounds (pending further kinome wide cross screening)

Table 8 – List of gap kinases – kinases that we currently do not cover with a compound that has sufficient selectivity and/or potency

